# Machine learning goes wild: Using data from captive individuals to infer wildlife behaviour

**DOI:** 10.1101/2019.12.18.881011

**Authors:** W. Rast, S. E. Kimmig, L. Giese, A. Berger

## Abstract

1. Remotely tracking distinct behaviours of animals using acceleration data and machine learning has been carried out successfully in several species in captive settings. In order to study the ecology of animals in natural habitats, such behaviour classification models need to be transferred to wild individuals. However, at present the development of those models usually requires direct observation of the target animals.
2. The goal of this study was to infer behaviour of wild, free roaming animals from acceleration data by training behaviour classification models on captive individuals, without the necessity to observe their wild conspecifics. We further sought to develop methods to validate the credibility of the resulting behaviour extrapolations.
3. We trained two machine learning algorithms proposed by the literature, Random Forest (RF) and Support Vector Machine (SVM), on data from captive red foxes (*Vulpes vulpes*) and later applied them to data from wild foxes. We also tested a new advance for behaviour classification, by applying a moving window to an Artificial Neural Network (ANN). Finally, we investigated four strategies to validate our classification output.
4. While all three machine learning algorithms performed well under training conditions, the established methods, RF and SVM, failed in classifying distinct behaviours when transferred from captive to wild foxes. Behaviour classification with the ANN and a moving window, in contrast, inferred distinct behaviours and showed consistent results for most individuals.
5. Our approach is a substantial improvement over the methods previously proposed in the literature as it generated plausible results for wild fox behaviour. We were able to infer the behaviour of wild animals that have never been observed in the wild and to further illustrate the outputs credibility. This framework is not restricted to foxes but can be applied to infer the behaviour of many other species and thus empowers new advances in behavioural ecology.

## INTRODUCTION

Animal-borne sensors such as temperature loggers, salinity loggers or microphones are used to study a wide variety of parameters in wild animals without disturbance by human observers (review in Cooke et al., 2004). In the study of movement ecology of species (Nathan, 2008), animal borne sensors make it possible to track the positions of wild animals. The first attempts to remotely track animal locations were made in the 1960s through VHF telemetry (Craighead and Craighead, 1972). In more recent years it has become common practice to track animal locations with satellite systems (Hebblewhite and Haydon, 2010), allowing us to study where individuals dwell. However, the spectrum of ecological questions that can be addressed by using location data alone is limited. By combining such data with behavioural data, more in-depth studies of species will become possible (Nathan et al., 2012). Yet, in contrast to recording locations, remotely tracking the behaviour of free-ranging animals is not well established at this point.

The principal underlying remote-tracking of behaviour is to attach accelerometers to animals to record their body movement. The first major study utilizing acceleration data to study the behaviour of animals was conducted in 1996 (Yoda et al., 1999). Since then, many studies have shown that acceleration data can be used to infer the behaviour of animals by employing various machine learning algorithms (Nathan et al., 2012; Bidder et al., 2014). To train these algorithms for pattern recognition and data classification, the acquisition of acceleration data was coupled with direct observation of the behaviours of the tagged animals. Using one portion of this ground-truthed data set to train the algorithm and another portion to infer behaviour from it is allows validation of the inferred behaviour.

Extrapolating behaviours from acceleration data of wild individuals is a challenge, since there is no possibility for direct output validation. Some models were trained and validated on the same wild individuals (Yoda et al., 2001; Tsuda et al., 2006; Studd et al., 2019), which requires direct observation of the studied individuals at least for a certain period of time. However, the promising advance of behaviour classification through machine learning is the ability to study the behaviour of wild animals without observing (and possibly disturbing) them. Furthermore, direct observation may often not be a feasible option, especially when target species are elusive or cryptic.

For other models, additional sensors such as GPS (Williams et al., 2014) or depth and speed sensors for aquatic species (Yoda et al., 2001; Tsuda et al., 2006; Zimmer et al., 2011) were employed to identify the behaviours executed. In these cases, the information from the additional sensors was used to investigate the behavioural context the animal was in at the time of data recording in order to delimit likely behaviours. For studies in which no validation was possible, various behaviours were grouped into broad, easily distinguishable categories to reduce confusion of similar behaviours (Halsey and White, 2010; Grünewälder et al., 2012). Thus, accurately inferring distinct behaviours of wild individuals still poses a problem.

The Random Forest (RF) and the Support Vector Machine (SVM) are popular approaches to infer animal behaviour from acceleration data and have yielded good results under training conditions (Nathan et al., 2012; Tatler et al., 2018). However, we are not aware of any study successfully transferring a behaviour classification model trained on captive individuals to wild individuals.

In this study we train three different machine learning algorithms on acceleration data from captive red foxes (*Vulpes vulpes*) and apply them to data from wild individuals, to infer wild fox behaviour. We address the limitations of the two above-mentioned machine learning algorithms in inferring behaviour from wildlife data and provide a new framework using a moving window and an Artificial Neural Network (ANN). Furthermore, we address the issue of how to validate the inferred behaviour when observation of free-ranging individuals is not a feasible option. We propose four strategies to assess the credibility of the output by combining the classified behaviour with GPS and temporal information.

With both captive and wild individuals tagged with the same type of GPS-acceleration sensors, captive individuals were observed to train the models and wild individuals were used to apply them. This set-up together with our novel approach enables us to improve the use of machine learning for behaviour classification. We thus hope to empower behaviour classification of wildlife through acceleration data in the future.

## MATERIAL AND METHODS

### Data collection and acceleration logger setup

Animal catching and handling have been approved from the State Office for Health and Social Affairs, department of veterinary affairs (permit number: IC113-G0211/15) and the ethics committee of the Leibniz Institute for Zoo and Wildlife Research in Berlin (permit number: 2015-03-04) and have been conducted according to applicable national and international guidelines. Approvals have been received prior to beginning research. To reduce stress during handling, all foxes got anesthetized before the deployment of radio collars. For anaesthesia we first used a long established mixture of Xylazin (10-16mg/kg) and Ketamin (12-20mg/kg) and later switched to an improved mixture of Ketamin (4mg/kg), Medetomidin (70µg/kg) and Midazolam (0,6mg/kg) that is better tolerated.

For gathering the training data set, we deployed UHF-GPS collars (E-obs GmbH, Munich, Germany) on two female red foxes in a game park enclosure. For the field dataset, data from 9 wild foxes was used that were radio-collared by Kimmig et al. in the city of Berlin, Germany between 2015 and 2018 using the same tag type and settings as used for the captive individuals.

The acceleration loggers that were embedded in the UHF-GPS collars were set up to measure acceleration every two minutes. Data was recorded for three axes perpendicular to each other at a sampling rate of 33.33Hz per axis. For every high-resolution recording interval, the so-called burst, 110 measurements per axis were made, which resulted in intervals of 3.3 seconds.

To train the algorithms we used the raw ground-truthed data of the captive foxes that were observed during the recording of acceleration data. A specific pinger signal indicated the start of each burst. This could be detected acoustically with an UHF Wide Range Receiver that was set to the unique frequency of the collars (see Giese 2016). During a burst the animal in focus was observed closely and the behaviour was noted. Each observation was linked to the corresponding acceleration burst via the unique timestamp.

Due to slight shifts in the collars’ timestamps, the raw acceleration data of a number of consecutive bursts - ideally encompassing a distinctive change in behaviour (e.g. resting followed by trotting) - was visually inspected and compared to the noted behaviours. The timestamps of observations were corrected accordingly.

After excluding all bursts containing more than one behaviour, 4159 bursts of six different behaviour classes were used as the input for the model training (feeding: 367, grooming: 1140, resting: 2114, caching (bury food to consume it later): 197, trotting: 179, walking: 162; see ethogram in Table S1).

### Data preparation

We calculated summary statistics from the raw acceleration data, separately for each burst, to serve as predictors for the machine learning algorithms. The following predictors were calculated per axis: mean, standard deviation, inverse coefficient of variation, variance, skewness and kurtosis. Additional predictors represent combinations of all three axes and were calculated according to the corresponding literature: q (Nathan et al., 2012), pitch and roll (Collins et al., 2015) and overall dynamic body acceleration (ODBA) (Wilson et al., 2006). For the Artificial neural network, we added the whole spectrum of a Fast Fourier Transformation of every axis to the set of predictors. For a complete list of predictors see Table S2. We performed all data transformations and the construction of the ANN in R (R Core Team, 2018) and Rstudio (RStudio Team, 2016). The sum_data function in the accelerateR package <u>(W. Rast, unpublished data.)</u> was used for summary statistic and Fast Fourier Transformation calculation.

### Data classification

#### Established methods: Support Vector Machines (SVM) and Random Forest (RF)

Support Vector Machines separate data of different classes from each other by constructing a hyperplane between them. Classification of new data is subsequently based on their relative position to the hyperplane. By default, the classification is binary. For applications with multiple classes, more hyper-planes between classes will be constructed (Cortes and Vapnik, 1995). We used the implementation of an SVM in the R package “e1071” (Meyer et al., 2017) with the kernel type “radial”.

Random Forests are an improvement of the classical Classification and Regression Trees (CART) (Breiman et al., 1984). While in CART all predictors are used, the RF picks a random subset of predictors to fit a tree. This is repeated several times, and the final prediction is the result of all trees combined by a majority rule (Breiman, 2001). We used the implementation of an RF in the R package “randomForest” (Liaw and Wiener, 2002) with the standard settings using 500 trees.

#### Artificial Neural Network (ANN)

ANNs are similar to biological neural networks and consist of multiple nodes that are distributed over several layers and interconnected (Jain et al., 1996). Nodes are activated based on the input variables (predictors) and an activation function. In the simplest cases, this function is a summation of all input variables that are passed to a specific node. These functions also include weights that change the influence of every input variable and are set during the training phase. For training, a ground-truthed data set is needed on which the ANN establishes the node connections and the weights so that the output of the ANN corresponds to the target classes of the model data. The activation or non-activation of nodes serves as input for the next layer of nodes. The last layer usually consists of nodes representing the target classes. Their activation leads to the assignment of data to a class.

For our study, we chose a three-layer network with the output of the last layer being a specific behaviour. We used a feed-forward type architecture for the ANN and used the keras package (Allaire and Chollet, n.d.) to implement it.

### Moving window

One strategy that has been tested with continuously recorded data is to apply a moving window to partition the acceleration data and to compute summary statistics for each of the resulting segments. In different studies, these windows could partially overlap or not overlap at all (Hokkanen et al., 2011; Lush et al., 2016; le Roux et al., 2017). An application example very similar to our approach is the assessment of car driver aggressiveness using continuous data by Ferreira et al. (2017). However, to our knowledge, this approach has never been used on burst data in wildlife ecology.

We applied a moving window to every recorded burst to increase the sample size of our data set, since it was found that ANNs show better performance with increasing sample size (Nam and Schaefer, 1995; Zhang et al., 1998) and require large training data sets (Srivastava et al., 2014). In the first set, this window reduced the amount of data within the burst from the original 110 measurements down to a subset of the window length. We then computed the summary statistics and Fast Fourier Transformation (Table S2) for this subset. In a second step, the window was moved by one position so that the first measurement of every axis was removed and one new measurement for every axis was added to the end of the window. We then computed all variables for the second window and so forth. The window was moved until it included the last measurement of the original burst, resulting in a number of predictor sets representing the same burst. In contrast to extracting random subsets, this approach preserves the order of the data for a specific behaviour, and we were able to calculate the Fourier spectrum which is dependent on the correct order of measurements.

### Model evaluation

To evaluate the model performance of all three machine learning algorithms, we trained them on 70% of the data (training data). We then inferred the behaviour for the remaining 30% (test data) by classifying them with the trained model and assigning a specific behaviour to each burst accordingly (or assigning “other”, respectively as described below). Since the number of observations per behaviour class differed, we split the data so that the original proportions of behaviour counts were similar in the training and test data sets. We applied the moving window to the training and test sets after the split. We calculated the recall (true positives / (true positives + false negatives)) and precision (true positives / (true positives + false positives)) (Bidder et al., 2014) for each behaviour.

To reduce confusion of behaviours, a threshold was set for the ANN assignments. Only behaviour assignments that exceeded a probability of 0.7 were accepted. All assignments below that threshold were classified as “other” behaviour. This was necessary to account for the fact that captive individuals may not execute the full range of behaviours available to the species, which would lead to some behaviour (e.g. hunting or fighting) not being included in the model. If wild individuals displayed any of these behaviours, they could be incorrectly classified as one of the behaviours included in the model. We expect that such classifications would be assigned at low probability, so that we can avoid these errors by implementing the threshold. Similarly, recordings in which the individual changed its behaviour during a burst should not be characteristic for any specific behaviour and therefore should also fall below the threshold.

### Model selection

Artificial Neural Networks are used for a variety of tasks such as image recognition, sentiment analysis or regression. The necessary sample size and ANN architecture depend on the specific task (Allaire and Chollet, 2018). Finding the optimal properties for the best performing ANN is not achieved by a scientific method but rather by trial and error (Zhang et al., 1998). To find the best window size we trained the ANN on window sizes from 20 to the full 110 and finally decided on 79. We evaluated all models by calculating the recall, precision and the proportion of “other” behaviours. As recall and precision are calculated for each behaviour, we first computed their means and then calculated the mean of the resulting mean recall and precision as well as the proportion of “other” behaviours. The latter was subtracted from one to be on the same scale as recall and precision. A General Additive Model (GAM) was applied to the calculated means for all window sizes. We calculated the slope m of the GAM fit for each window size using the difference quotient m=(Δy_n_-Δy_n-1_)/(Δx_n_-Δx_n-1_). Variable x corresponds to the window size and y to the calculated model performance, n corresponds to a specific window size and n-1 to the previous window size. A window size of 79 provided the best trade-off between small window size and high performance (see “Model selection” in the Results).

### Application to wild individuals

For our subsequent analysis of behaviour inference, we selected wild foxes for which at least three consecutive months of acceleration data were available (N=9). We considered all months in which data was recorded for at least half of the month. In addition to the acceleration data, the tags recorded GPS positions every four minutes for the first eight weeks, after that every 20 minutes (GPS for fox “Gerlinde” was only recorded every 20 minutes). Using acceleration informed GPS measurement, this interval was reduced to every four hours when a fox was inactive. We trained all three classification models on the complete ground-truthed dataset of the captive foxes and applied the trained model to classify the data of the wild foxes. As the moving window results in multiple behaviour outputs for each burst, only one behaviour was assigned to each burst, following majority rule.

### Validation of behaviour assignments

We assessed the plausibility of the ANNs’ behavioural assignments by examining the following four aspects: (i) biological credibility of the behaviour assignments (ii) consistency over individuals and time (iii) coherence with the GPS data and (iv) coherence with ODBA.

To address biological credibility (i), we calculated the time-dependent composition of behaviours throughout the day and compared it to the literature on fox behaviour. As seasonal shifts can influence behavioural compositions, we separated the behaviour assignments by month. For each day within a single month, we counted the number of assignments of each behaviour (for each minute covered by the tag schedule) in the 24 hours. We further used the corresponding plots to (ii) visually compare the daily patterns over time and between individuals. (iii) We incorporated the given GPS information of the free-ranging individuals because we expected the GPS data to correspond with specific behavioural classes. For instance, spatial clustering of GPS data should correspond with stationary resting behaviour. We treated points as a cluster when consecutive GPS points were within a 50m radius of the first GPS point of that cluster. Points recorded more than 50m away were defined as the first point of a new cluster. Since it was possible that clusters consisted of only a single point, we only considered behaviour assignments to be spatially clustered when at least 10 classified behaviour items were assigned to the same cluster. We then calculated for each behaviour the proportion of behaviour assignments that were within a cluster. In addition, we investigated coherence of GPS based speed measure and movement related behaviour classifications (trotting and walking). We therefore calculated the speed the animal was moving at during acceleration measurement based on the spatial and temporal distances between consecutive GPS points. Due to independent schedules, GPS and acceleration data were not recorded exactly simultaneously. Hence acceleration data that was recorded within 10 seconds of a GPS measurement were considered. Finally, we (iv) compared the temporal distribution of ODBA values and behaviour assignments by constructing actograms using accelerateR.

## RESULTS

### Training conditions: Captive foxes

We could classify all six behaviours during the validation using SVM and RF. Classification success differed between the behaviour classes for both algorithms. We achieved the best classification success for resting and the lowest for caching and walking (Table 1). The confusion matrices (Table S3 and S4) showed that grooming and walking were confused more often compared to other behaviours. Overall, behaviours with a low recall also showed a low precision and those that had a high recall also showed a high precision. The exceptions were caching for the SVM with high recall (0.75) and lower precision (0.50) and walking for the RF with low recall (0.27) and higher precision (0.57). Recall can be interpreted as the proportion of behaviour events that were correctly classified. Trotting (SVM), for example had a recall of 0.72, meaning that 72% of all trotting events were correctly classified as trotting. Precision can be interpreted as the probability for an assignment to be correct. Trotting had a precision of 0.98, meaning that a single assignment of trotting is correct with a chance of 98%. Both algorithms show comparable results, with the most prominent differences in the recall of trotting and walking with an increase of 0.19 and a decrease of 0.32 respectively.

**Table 1:** Recall and precision of the classification output compared for support vector machine (SVM), random forest (RF) and artificial neural network (ANN). All algorithms are capable of classifying and inferring fox behaviour with a high success rate (exceptions are caching and walking for SVM and RF).

|  | **feeding** | **grooming** | **resting** | **caching** | **trotting** | **walking** |
| --- | --- | --- | --- | --- | --- | --- |
| **SVM** |  |  |  |  |  |  |
| recall | 0.65 | 0.80 | 0.96 | 0.75 | 0.72 | 0.59 |
| precision | 0.70 | 0.83 | 0.97 | 0.50 | 0.98 | 0.48 |
| **RF** |  |  |  |  |  |  |
| recall | 0.69 | 0.89 | 0.97 | 0.67 | 0.91 | 0.27 |
| precision | 0.78 | 0.81 | 0.97 | 0.61 | 0.96 | 0.57 |
| **ANN** |  |  |  |  |  |  |
| recall | 0.83 | 0.88 | 0.96 | 0.67 | 0.96 | 0.74 |
| precision | 0.84 | 0.95 | 0.98 | 0.83 | 0.91 | 0.71 |

Like RF and SVM, the ANN could predict all six behaviours during validation. Also, classification success differed between behaviour classes. However, classification success was higher for all behaviour classes except grooming and walking (recall for grooming decreased by 0.01 and precision for trotting decreased by 0.07). The most drastic change appears in the recall of walking with an increase of 0.47 (Table 1). The confusion of walking behaviour with grooming is reduced compared to the SVM and RF (Table S6). The proportion of assignments that did not surpass the threshold was 0.04.

Model performance of the ANN appears to be dependent on the window size (Fig. 1) and decreases towards both ends of the window size spectrum. Smaller window sizes seem to have a stronger impact on model performance than larger window sizes. The GAM fit has its maximum at window size 79, with the slope of the GAM fit close to 0. We thus considered 79 to be the best trade-off between model performance and window size and used it for the final model (see Discussion).

**Fig. 1:**
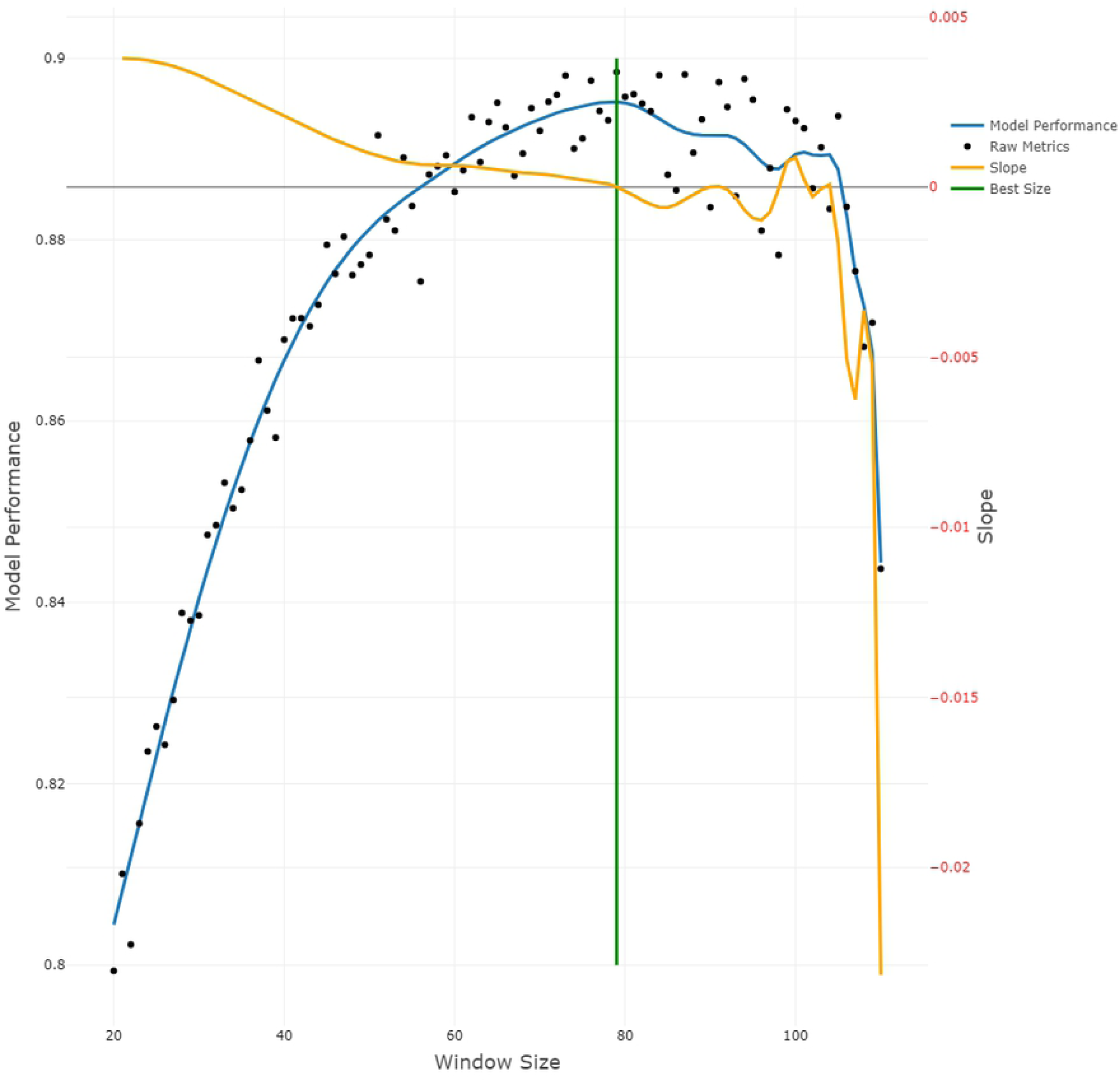
ANN model performance in relation to window size. Black dots show the computed performance values. The blue line is the result of a General Additive Model, k = 40 (Wood, 2017) fit. The y-axis on the left side labelled “Model Performance” corresponds to the Model Performance line (blue) and Raw Metrics points (black). The orange line is the calculated slope of the model performance, which corresponds to the y-axis on the right side labelled “Slope”. The green vertical line represents the best window size of 79.

### Field conditions: Application to wild foxes

We show here the detailed results for two wild foxes (“Que” and “Gerlinde”), whose collars yielded data over a whole year. For the seven remaining individuals, data is only available for shorter time periods. Plots and data for these animals are presented in the supplemental material.

When the trained SVM model was applied to classify the behaviour of the nine wild foxes, all bursts were classified as grooming. No resting, caching, feeding, trotting or walking events were detected. When using the trained RF model, most of the bursts were classified as grooming behaviour and the remaining ones as resting (Table 2, Table S5). When applying the trained ANN to the wild fox data, all six behaviour categories were assigned in all nine individuals. For the field data of “Que” and” Gerlinde”, a proportion of 1% did not exceed the 70% threshold and was therefore labelled “other”. For both foxes feeding, caching and walking were assigned at low rates (Table 2, Table S7 for the remaining foxes).

**Table 2:** Number of occurrences of every classified behaviour for the foxes “Que” and “Gerlinde”. Count and proportion of all behaviour assignments compared for support vector machine (SVM), random forest (RF) and artificial neural network (ANN).

| Individual | Measure | feeding | grooming | resting | caching | trotting | walking | other |
| --- | --- | --- | --- | --- | --- | --- | --- | --- |
| **SVM** |  |  |  |  |  |  |  |  |
| Que | count | 0 | 274792 | 0 | 0 | 0 | 0 | 0 |
|  | proportion | 0 | 1,000 | 0 | 0 | 0 | 0 | 0 |
| Gerlinde | count | 0 | 289248 | 0 | 0 | 0 | 0 | 0 |
|  | proportion | 0 | 1,000 | 0 | 0 | 0 | 0 | 0 |
| **RF** |  |  |  |  |  |  |  |  |
| Que | count | 0 | 212728 | 62064 | 0 | 0 | 0 | 0 |
|  | proportion | 0 | 0,774 | 0,226 | 0 | 0 | 0 | 0 |
| Gerlinde | count | 0 | 207576 | 81672 | 0 | 0 | 0 | 0 |
|  | proportion | 0 | 0,718 | 0,282 | 0 | 0 | 0 | 0 |
| **ANN** |  |  |  |  |  |  |  |  |
| Que | count | 1317 | 64092 | 166349 | 1875 | 32531 | 5749 | 2879 |
|  | proportion | 0.005 | 0.23 | 0.61 | 0.007 | 0.12 | 0.02 | 0.01 |
| Gerlinde | count | 2288 | 78194 | 171890 | 1016 | 30020 | 1664 | 4176 |
|  | proportion | 0.008 | 0.27 | 0.59 | 0.004 | 0.10 | 0.006 | 0.01 |

### Validation and credibility of the behaviour assignments

#### Biological credibility of the behaviour assignments (i) and consistency over individuals and time (ii)

Looking at the time-dependent composition of each individual’s behaviour (Fig. 2, Figs S1-S7), a similar pattern of behavioural composition over time is clearly noticeable (months without full data recording ought to be excluded for feasible interpretation). Clearly, there is a high proportion of resting behaviour during the middle of the day, while trotting is mostly inferred during dark hours. Trotting is also inferred more often than walking. There seems to be a seasonal change in resting behaviour, with resting events being more explicitly limited to the daytime in summer months. Feeding events are more often inferred during dark hours than during daytime, when mostly resting and some grooming are classified.

In the comparison between individuals some differences emerge. Some individuals, e.g., show less trotting (Fig. S2), more walking (Fig. S3) or much more grooming than others (Fig. S7). Despite this variation, the general pattern of behaviour composition appears very similar across all individuals.

**Fig. 2:**
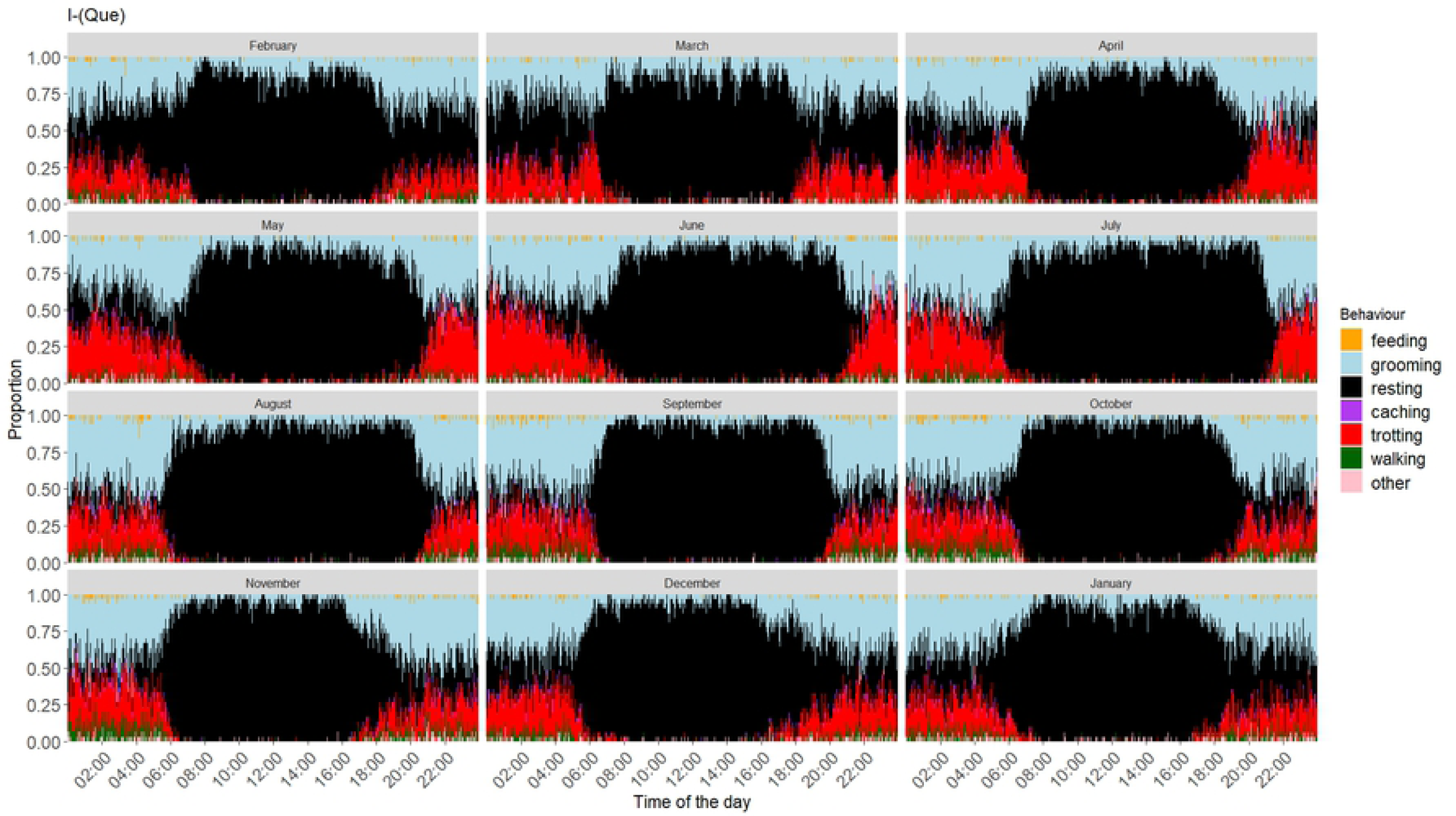
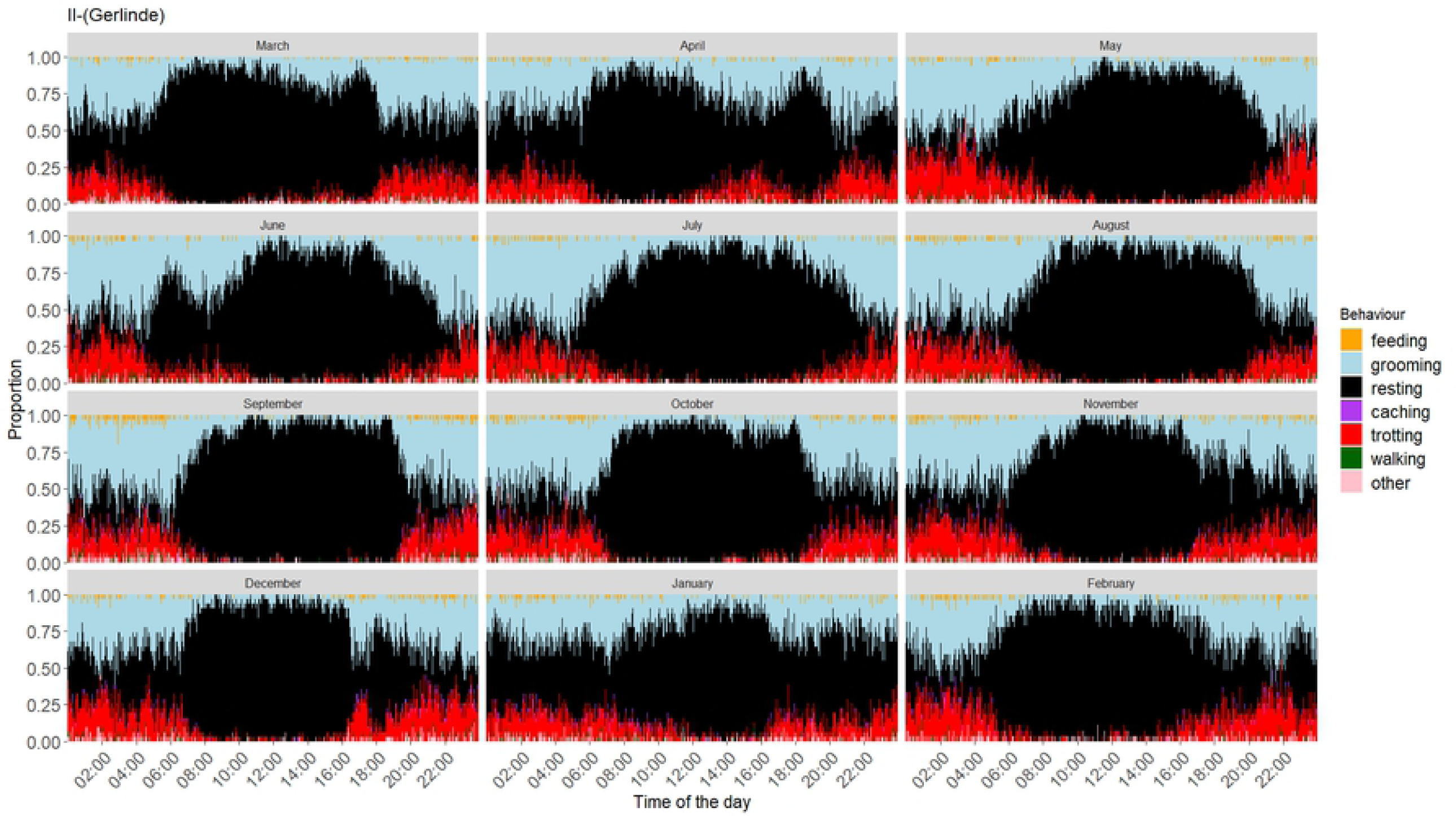
Time-dependent composition of behaviours of Que (I) and Gerlinde (II). Stacked bars represent the proportion of each behaviour at a given time of day, in each month. The data shown here span from February 2018 to January 2019 for Que and from March 2016 to February 2017 for Gerlinde.

#### (iii) Coherence with GPS

Resting behaviour appears to be highly associated with GPS clusters (Fig 3A), while all other behaviours are inferred mostly outside of clusters. This also applies to all other wild foxes (Figs S8A - S14A). For the analysis of the correspondence between behaviours and GPS based speed measurements, we used only acceleration data recorded within 10 seconds of a GPS fix (for Que: 7%; Gerlinde: 5%). Resting events were classified at lower GPS-based speed than trotting events (Fig 3B, Figs S8B - S14B).

**Fig. 3:**
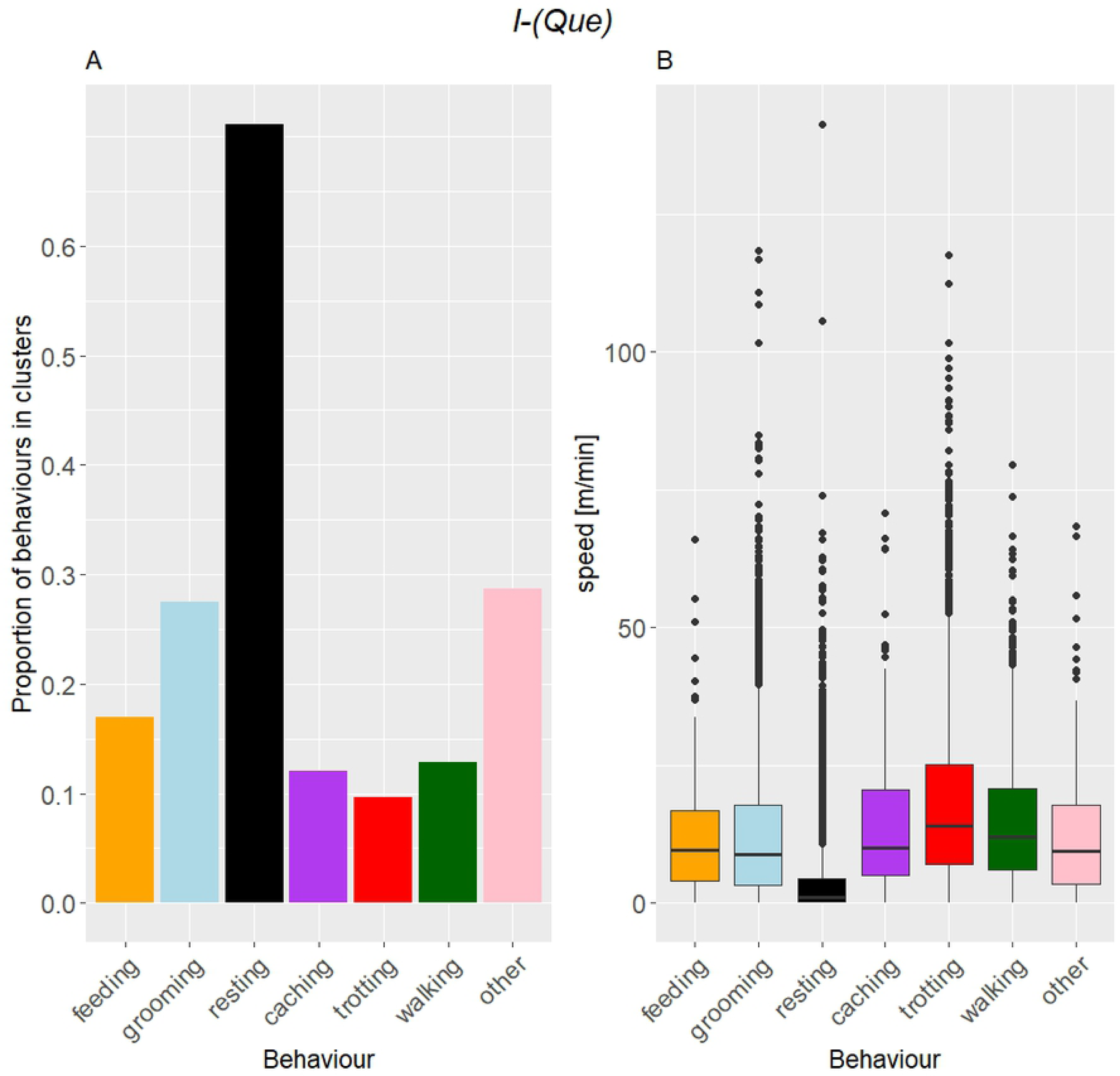
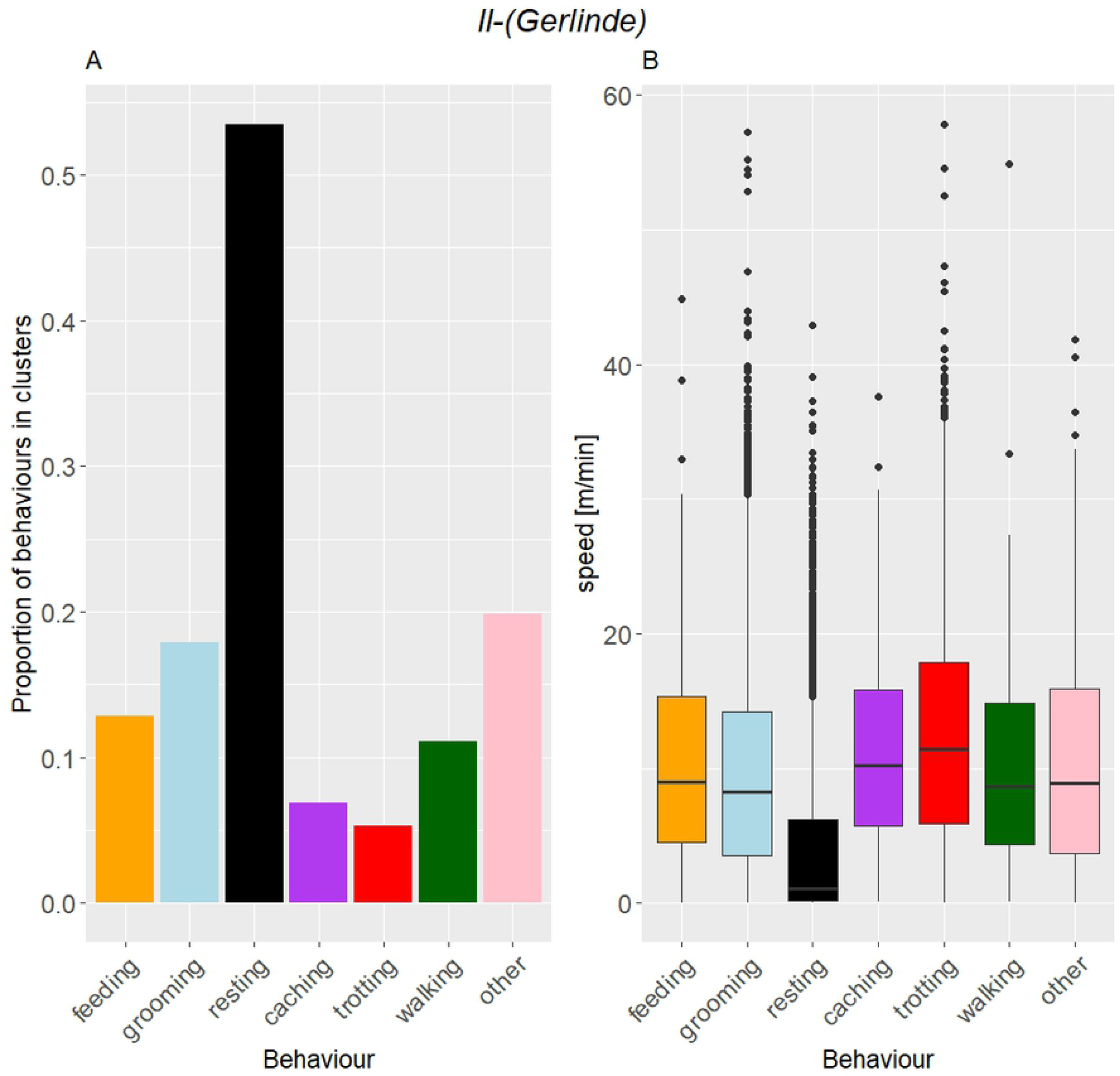
Behaviour assignments of Que (I) and Gerlinde (II) in relation to GPS clusters (A) and speed (B). (I) Resting shows the highest association with GPS clusters (71%) and trotting the lowest (9%). Resting events are associated with significantly lower speed than trotting events (Wilcoxon rank sum test, W = 3024826, p < 0.001). (II) Resting shows the highest association with GPS clusters (53%) and trotting the lowest (5%). Resting events are associated with significantly lower speed than trotting events (Wilcoxon rank sum test, W = 2286090, p < 0.001).

#### (iv) Coherence with ODBA

The actograms show that trotting is predominantly classified at times when ODBA values are high. Trotting, as well as high ODBA, occur mostly during night-time. Resting, in turn, is most often classified at times with low ODBA values (Fig. 4). This is also valid for all remaining foxes (Figs S15-S21).

**Fig. 4:**
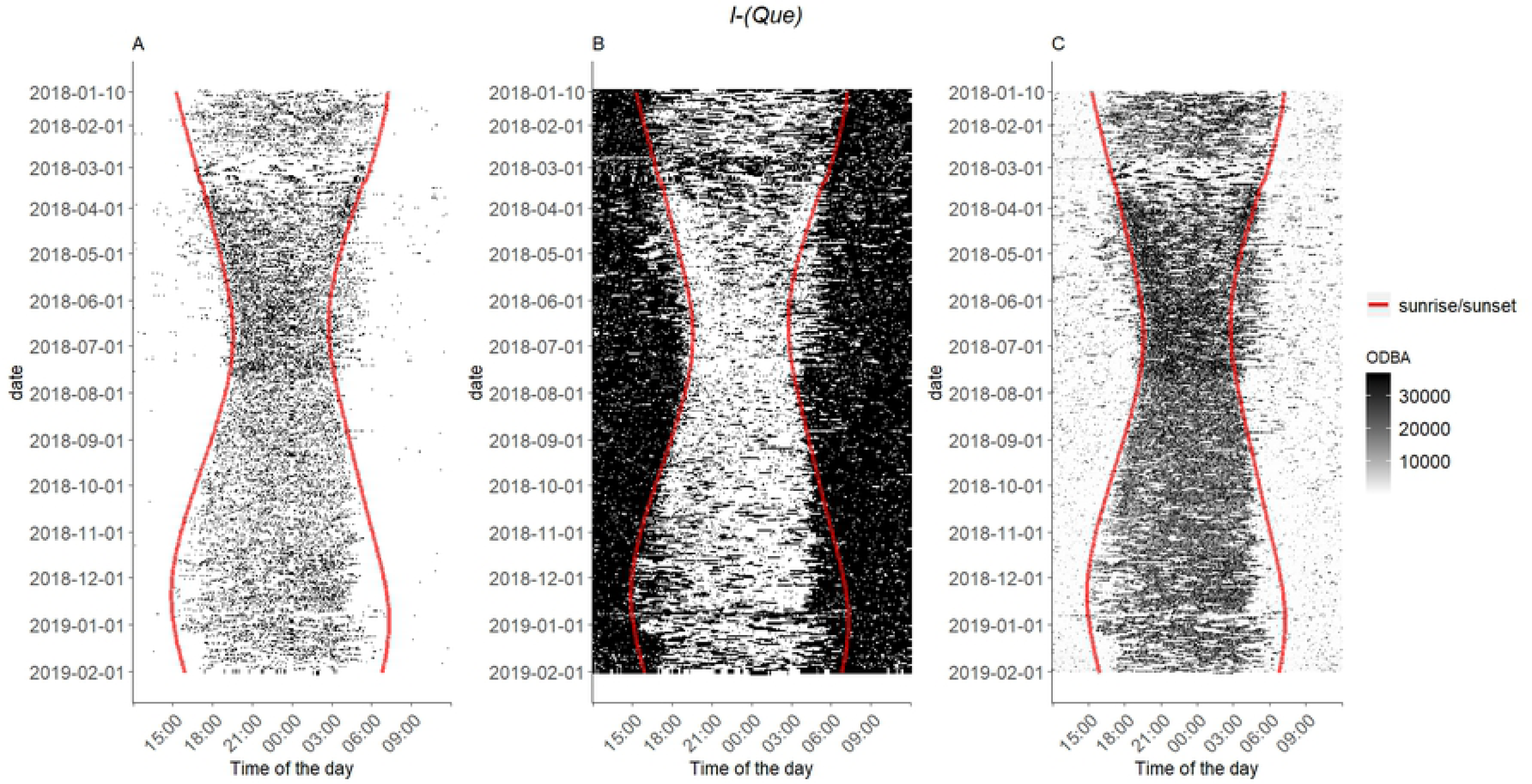
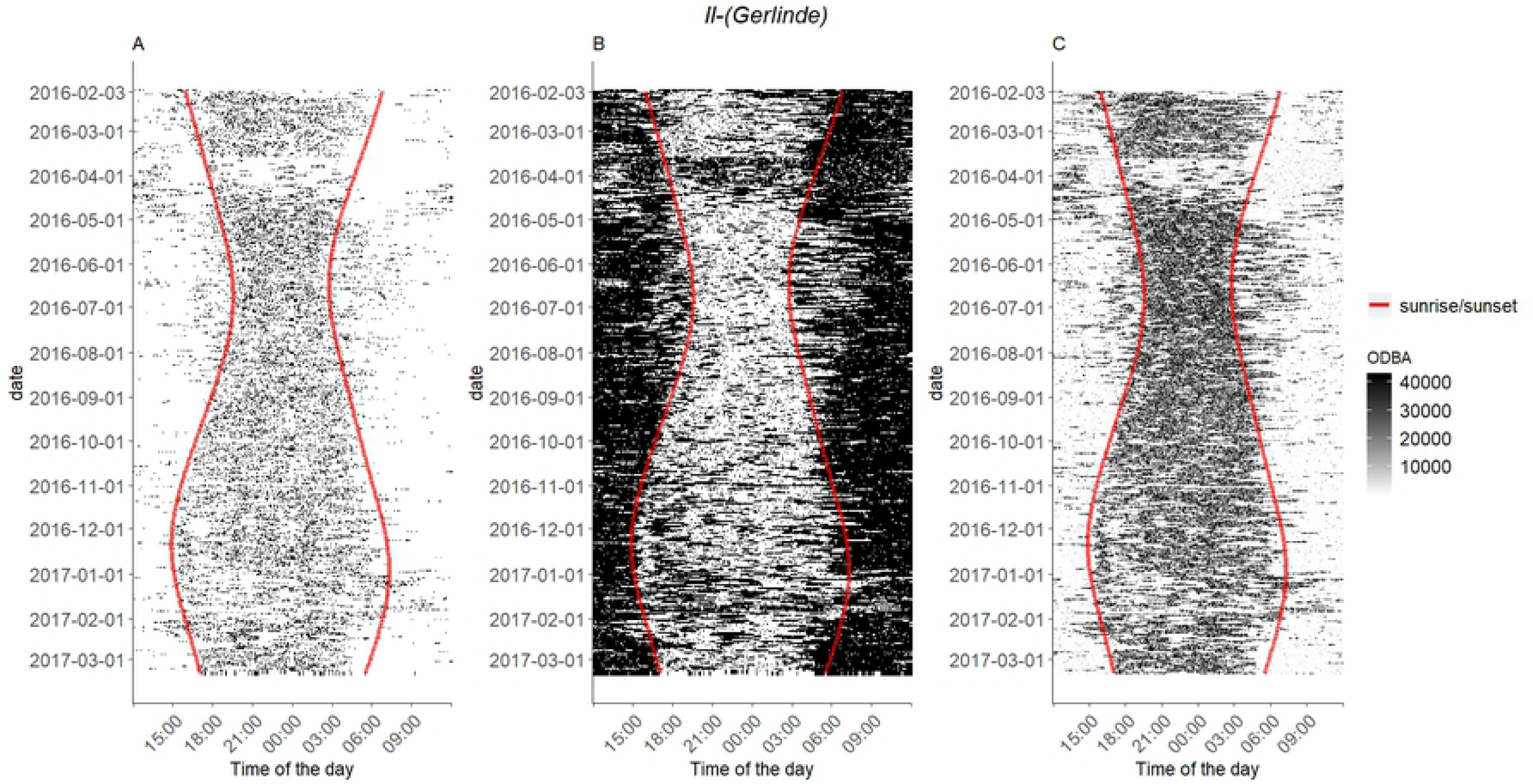
Temporal distribution of trotting (A), resting (B) and ODBA values (C) for Que (I) and Gerlinde (II). The red lines indicate sunset and sunrise. (A) Black spaces indicate times at which trotting behaviour was classified, whereas white spaces indicate classification of all other behaviours. (B) Black spaces indicate times at which resting behaviour was classified, whereas white spaces indicate classifications of all other behaviours. (C) Higher ODBA values are indicated by darker spaces.

## DISCUSSION

In the present study we sought to advance the abilities to remotely assess the ecology and behavior of animals in the wild without directly observing (and respectively disturbing) the target animals.We, therefore, tested the capacity of three machine learning algorithms (SVM, RF and ANN) to infer wild fox behaviour after training with a ground-truthed data set of two captive red foxes. Recall and precision were on similar levels for all three algorithms. The ANN with the moving window approach, however, was able to infer caching and walking behaviour much better than the other two. Both RF and SVM generally performed well in inferring behaviour during validation (Table 1) and showed comparable results to other studies (Nathan et al., 2012; Fehlmann et al., 2017; Kröschel et al., 2017). When applied to the wild foxes, however, they both failed to discriminate the different behaviours (Table 2, Table S5).

### Support vector maschine (SVM) and random forest (RF)

When applied to the data from wild foxes, SVM classified all behaviours as grooming, while RF distinguished only grooming and resting (RF). Considering that these datasets contain measurements for at least three months and that the GPS signal clearly showed the animals covering large distances, this classification is clearly unrealistic. In all cases, the models were trained and validated on measurements from the same captive individuals. Since both models correctly inferred behaviours of those captive individuals during validation, we suggest that SVM and RF thus failed to recognize similar patterns between wild and zoo-kept individuals because wild foxes showed characteristics in their behaviour that were too different from the captive foxes to be detected by those approaches.

This could be solved by training the model on ground-truthed data obtained from observing wild individuals that are logged (Grünewälder et al., 2012). However, as mentioned above, this is not always feasible, so it would be highly desirable to do trait classification without the necessity to directly observe wild animals. An additional problem of SVM and RF could be the recording of mixed behaviours. During model training, all recordings with mixed behaviours within one burst were excluded from the data set. However, it is a fair assumption that at least some recordings from wild individuals do contain more than one behaviour. Even though we implemented a probability threshold to account for these mixed bursts and - in the best case - classify them as “other” behaviour, these recordings might still pose an unclassifiable problem. Another issue is that grooming is classified to such a high proportion by both SVM and RF. As grooming behaviour is complex and may include different actions like licking, nibbling or scratching (see also Table S1), the resulting behaviour class probably includes a broad range of characteristic summary statistics and may thus be more easily confused with other behaviour classes.

### The artificial neural network (ANN)

The ANN showed comparable results to SVM and RF in model validation and consequently to the literature as cited above. However, when applying it to the same wild individuals the SVM and RF failed on, it classified a very different set of behaviours (Table 2). If we had only tested the two established approaches, we would have concluded that the transfer of a behaviour classification model trained on captive foxes to wild individuals is not possible. In contrast, the ANN with a moving window shows promising results that hopefully prompt further investigation into its potential use for wildlife ecology. The approach brings two additional advantages, besides the increased sample size: First, the likely better handling of recordings with mixed behaviours: By reducing the number of measurements per burst, the proportion of a potential second behaviour in the same burst is reduced. In some cases, this reduction may be enough to calculate similar summary statistics to a burst with only one of the behaviours. Second, the introduction of an ensemble learning effect: In case of a specific behaviour being performed in an unusual way or a burst containing more than one behaviour, it will be harder to infer the correct behaviour. As the moving window creates 32 subsamples of the original burst, using a majority vote for the resulting 32 assigned behaviours can reduce the uncertainty of the assignments.

The best size for the moving window was determined based on the maximal performance and slope of the GAM fit in the simulation plot (Fig. 1). Our aim was to find a window size with a high mean performance that is small enough for generating sufficient data. Unfortunately, reducing the window size was found to negatively impact model performance (Tatler et al., 2018). Performance seemed to be at its maximum at window size 79. Larger windows would result in fewer subsets at worse performance, while performance also decreased for window sizes smaller than 79. Considering the slope of the GAM fit, the performance changed only marginally at window sizes 78 and 80 compared to 79. Therefore, we expect the model to perform similarly well at these window sizes. We suggest considering the slope because this approach may not always show a clear maximum like in our case. In cases of multiple maxima or plateau formations, the slope will help to inform on the smallest window size with the best performance.

A problem that may occur with the moving window approach is incorrect classification through overfitting (Dietterich, 1995). By creating several similar subsets of the same burst, the ANN could build a model that fits the training data too well, i.e. even slight differences between training data and new data of the same behaviour class would result in the classification of different behaviours, with an overfitted model. Variation within the behaviour classes could also cause incorrect classifications if a single behaviour is realised outside the normal variation. There is no clear method to distinguish between these two causes of incorrect classifications. However, in the following we discuss the credibility of the assigned wild fox behaviour and argue that the moving window approach does not introduce overfitting.

### Output credibility

The classification results of the classic methods appear to be obviously incorrect. At first glance, the output of the ANN appears more plausible than the output of RF and SVM, on account of all six behaviours getting classified. Still, the actual accuracy cannot be determined as wild individuals could not be observed. Since this may be true for most tagged wildlife, we provide four strategies to indirectly assess the credibility of the ANN output.

When looking at the time-dependent composition of behaviours, they appear quite consistent over individuals and time. Generally, some variation between individuals is apparent and some behaviour events seem to be misclassified. Individual differences in moving behaviour, for example, may result in the assignment of either walking or trotting when the algorithm is not accurate enough. However, an overall pattern is evident for most of the foxes and the temporal distribution appears plausible: The ANN output suggest that the foxes predominantly rest during the day and are active at night as well as during twilight (Fig. 2, Figs S1-S8) which corresponds well with described nocturnal-crepuscular activity patterns of red foxes (Adkins and Stott, 1998; Díaz-Ruiz et al., 2016).

There also seem to be seasonal changes in these patterns, with fewer resting events during dark hours in the summer months. Although there are only two complete year-round datasets available, this pattern appears reasonable as the nights in summertime at this longitude are much shorter than during winter months (in Berlin, the daily dark period ranges from 7 to 17 hours). Thus, foxes should use the full night-time spectrum in summer for their activities, while during winter the higher availability of potential activity time allows nocturnal resting events. Like trotting and walking, feeding is mostly classified at night-time. Feeding events occur in no clustered manner. The mixture of movement and feeding events reflects the feeding ecology of foxes which do not feed on large prey. Foxes mostly hunt for small prey like mice and voles and often rather scavenge than hunt, especially in urbanized areas (Doncaster et al., 1990; Contesse et al., 2004).

We also used GPS data to relate the occurrence of GPS-clusters as well as GPS-based speed values to the assigned behaviour classes. In particular, we focused on resting and movement behaviour (trotting and walking), with an obvious connection to be expected. Resting as a stationary behaviour should get classified predominantly at locations where GPS points are clustered (Fig. 3). We found that 38% to 74% of resting bursts were located within such a cluster. The remaining bursts may reflect cases were the individual had just temporarily stopped moving. Standing still or sitting briefly during an active phase would also be classified as resting but may not be associated with a GPS-cluster.

Trotting as a locomotive behaviour was expected to show low association with GPS-clusters and was only classified at a cluster for 2% to 9% of all bursts (Fig. 3, Figs S8-S14.). As foxes move away from or to a resting site, it is reasonable for some trotting to be classified within GPS clusters. The analysis did not target feeding, caching or grooming, as those behaviours can be performed in a clustered or non-clustered manner. The analysis of behaviour assignments in relation to speed shows a reverse picture: Behaviours that show a weak association with GPS clusters show a higher speed and vice versa (Figs S8-S14). However, we could use only 5% to 15% of all data for the speed analysis (Table S8). The method described here is hence more applicable when the recording of location and acceleration data is better synchronised.

Finally, we analysed the ODBA, an indicator of body movement (Wilson et al., 2006) that was shown to correspond well with the activity level of specific behaviours (Tatler et al., 2018). When we compared the temporal distribution of ODBA values to that of the classified trotting and resting events, we saw an association of high ODBA values and trotting behaviour and low ODBA values and resting behaviour, respectively. Again, the nocturnal-crepuscular activity pattern was visible (Fig. 4, Figs S15-S21).

While the abovementioned examples appear conceivable, the interpretation of some behaviours may be puzzling, and their biological credibility is difficult to gauge. For instance, we could not identify any pattern for caching behaviour, and grooming seems to be generally over-classified. Its complexity and the resulting variability in the training data set may increase misclassification of unknown behaviours, especially when considering that six behaviours do not represent the full variety of behaviour that this mobile carnivore displays in the wild. A broader training dataset of more captive individuals could possibly improve the output of the ANN and permit more precise recognition of specific behaviours.

However, our results suggest that the behaviour inferred by the ANN corresponds well with the actual behaviour of the logged foxes. Despite some unsolved issues, the ANN thus seems to be a promising approach to infer wildlife behaviour, even in cases where methods suggested by existing literature fail.

## Conclusion

We provide a framework to use acceleration data and an Artificial Neural Network to infer the behaviour of wild foxes, using a training data set obtained from captive individuals. We also present four strategies to address the plausibility of such behaviour classification output when no direct validation is possible. Although not all validation strategies may be applicable for every species, this framework should not be restricted to the studied species. The successful application of the ANN for behavioural classification on field data offers exciting potential to study the behaviour of animals in the wild without direct observation.

## Author Contributions

AB, SEK and WR contributed to the design of this research. LG and SEK performed the field work. WR conducted data analysis. SEK and WR wrote the manuscript. All authors contributed critically to the drafts and gave final approval for publication.

## Data Accessibility Statement

Raw data of the captive foxes from which the machine learning model was build will and R code will be made available on Dryad upon publication. The data of the wild foxes is stored on Movebank and can be shared upon request.

## Acknowledgements

We are indebted to the Stiftung Naturschutz Berlin for providing funding for the project. We would like to thank Masahiro Ryo for suggesting the use of moving windows and Konstantin Börner, Janina Radwainsky and Frank Goeritz for their contribution to field work. Finally, we would like to thank Miriam Brandt for her constructive criticism to improve this manuscript.

